# Integration-coupled activation of promoterless combinatorial pathway libraries in *Clostridium* avoids burden during DNA assembly

**DOI:** 10.64898/2026.01.20.700586

**Authors:** Pawel M. Mordaka, James Williamson, John T. Heap

## Abstract

Combinatorial DNA design and assembly is an efficient and pragmatic way to obtain high-performing metabolic pathway designs quickly. However, implementation may require organism-specific technical barriers to be overcome. Firstly, suitable expression control parts such as promoters and ribosome-binding sites (RBSs) which provide a suitable range of expression levels need to be identified or developed. Secondly, these need to be assembled into pathway-encoding combinatorial libraries of sufficient size, quality and diversity. For organisms with transformation frequencies too low to allow direct transformation of library assembly reactions, such as many *Clostridium* spp., assembly and amplification is typically carried out using *Escherichia coli*. However, if constructs are deleterious (or ‘burdensome’) to *E. coli*, which is often the case when using *Clostridium* genetic parts, poor libraries may be obtained. Here we develop a new approach called integration-coupled activation of promoterless sequences (ICAPS) to overcome this barrier and therefore enable combinatorial assembly in *Clostridium*. Libraries were designed and assembled as promoterless synthetic operons, preventing expression during DNA assembly, and expression was only activated later, when constructs were integrated into the host genome downstream of a promoter. Variation of expression levels was achieved using a range of context-resistant RBS sequences. This approach was used to produce a *Clostridium acetobutylicum* library with combinatorial expression variants of an introduced hexanol pathway. This proof of concept provides a generally-applicable approach to implement combinatorial metabolic pathway-encoding libraries in *Clostridium* spp., circumventing the excessive strength of *Clostridium* expression control parts in *E. coli*, and is applicable to other organisms.

## Introduction

To transition the global economy towards sustainable biobased production, a range of organisms with different properties will be required to fulfil a range of bioproduction niches. *Clostridium* spp. were used widely during the early 20th century for the industrial production of acetone, ethanol and butanol, but bioproduction was later replaced in favour of cheaper production from fossil carbon [1]. Recently, *Clostridium* spp. have been undergoing a renaissance, building on historic examples that were proven feasible at scale with the application of modern bioprocess engineering and strain engineering.

As well as their natural production of useful products, the ability of *Clostridium* spp. to utilise a range of carbon sources is of interest, including industrial waste products such as lignocellulose, CO and CO_2_. These feedstocks do not compete with food production and are often available at low or potentially even negative cost (in the case of wastes with a disposal cost) making them ideal for the production of low-value, high-volume chemicals such as solvents [2]. LanzaTech and others are developing *Clostridium*-based fermentation technologies that can be retrofitted to high carbon-emitting industries, such as steel works, and also developing the use of gasification of biomass as a feedstock for fermentation [3]. The prospects and challenges of using *Clostridium* spp. at industrial scale are well reviewed by Vees *et al* [2].

As well as a range of organisms, the bioeconomy also requires the ability to build, test and optimise metabolic pathways in these organisms. *Clostridium* spp. were historically limited by a lack of tools for strain engineering, but are now relatively well served by tools including modular shuttle vectors [4], genetic parts such as promoters and selection markers (reviewed by [5]), directed genomic modification methods such as the bacterial Group II intron-based ClosTron system [6,7], integration using ‘ACE’ asymmetric homologous recombination [8] and examples of CRISPR in species including *Clostridium beijerinckii* [9], *Clostridium cellulolyticum* [10] and *Clostridium ljungdahlii* [11]. These foundational genetic tools for *Clostridium* spp. enable work towards implementation of more advanced approaches, such as highly-parallel combinatorial assembly.

Combinatorial DNA design and construction is an efficient and pragmatic way to achieve high-performing metabolic pathway designs quickly, leading to recombinant strains which produce desired products effectively, without prior knowledge of the optimal expression levels of individual coding sequences (CDSs). This is achieved by building libraries of metabolic pathway-encoding construct variants in which enzyme CDSs are placed under the control of a varied set of expression control parts (in bacteria mainly promoters, ribosome binding sites (RBSs) and terminators) to sample different designs in a ‘design space’. Golden Gate type multi-part DNA assembly systems using a standard framework, such as Start-Stop Assembly [12,13], are ideal for design and assembly of such libraries, which may use monocistronic, operon or hybrid designs. In Start-Stop Assembly (and other systems), any chosen mixture of parts in the appropriate format, at any chosen stoichiometry among the parts, can be used instead of an individual part at any position. The choice represents a trade-off between library diversity and screening practicality. We usually begin with standard mixtures of six promoters and six RBSs [12] which have proven effective in many cases. These libraries can be screened for variants that exhibit the desired traits, typically high yield and productivity, caused by combinations of enzyme expression levels that provide high flux towards the product, cause minimal accumulation of potentially toxic by-products or intermediates, avoid excessive protein expression, and show good cell growth [14]. These potential negative effects of heterologous expression have been formalised and studied as ‘metabolic burden’ [15,16]. Combinatorial methods have been used for functional optimisation of gene clusters [17] and improving production of a variety of products, including carotenoids, terpenoids, violacein, riboflavin, 2,3-butanediol and fatty acids (reviewed in [14,18]).

In this work we aimed to establish an approach allowing the construction of combinatorial metabolic pathway libraries in *Clostridium* spp. As an example to develop and test this approach, we constructed a hexanol production library in *Clostridium acetobutylicum*, building upon the native CoA-dependent branched fermentation pathway in this organism (**Figure 4**). Extension of the native clostridial 1-butanol pathway requires expression of an additional thiolase, such as β-ketothiolase Bktb from *Cupriavidus necator*, to condense butyryl-CoA and acetyl-CoA to 3-hydroxyhexanoyl-CoA. It is not known whether the native acyl-CoA dehydrogenase Bcd is able to reduce hexenoyl-CoA to hexanoyl-CoA. It has been shown that this step can be catalysed in *E. coli* by heterologous NADH-dependent trans-enoyl-CoA reductase Ter from *Treponema denticola* [19]. However, overexpression of the latter enzyme in *C. acetobutylicum* from a strong *ptb* (phosphotransbutyrylase) gene promoter resulted in a decreased butanol/acetate ratio [20]. Moreover, to improve alcohol titers, overexpression of the bifunctional aldehyde/alcohol dehydrogenase may be required. Bifunctional alcohol/aldehyde dehydrogenase AdhE2, used previously in a recombinant pathway in the *E. coli*, was shown to reduce hexanoyl-CoA to 1-hexanol, but it also showed activity towards acetyl-CoA and butyryl-CoA [19], which are intermediates in the pathway. Therefore, fine tuning of Bktb, Ter and AdhE2 expression levels is required to generate a strain with a functional 1-hexanol pathway with flux towards the final product rather than ethanol or butanol, making this an interesting and challenging case for the design or combinatorial development of a metabolic pathway-encoding construct (**Figure 4**). Previous work on hexanol production in *Clostridium* spp. has focused on fermentation conditions of species already capable of producing hexanol natively, such as *Clostridium carboxidivorans*, including in co-culture with *Clostridium kluyveri* [21–23]. Pathways from known hexanol producers have also been integrated into other *Clostridium* spp. [24].

Here we show how the excessive strengths of *Clostridium* promoters in *E. coli* causes a barrier to combinatorial assembly, we develop a new general approach to overcome this barrier, and we validate the approach using a hexanol pathway as a proof of concept. This work provides methodological foundations to apply combinatorial assembly for improved bioproduction in *Clostridium* and other organisms.

## Results

### Synthetic Clostridium promoters are too strong in E. coli, preventing standard combinatorial assembly

Initially, we set out to use the standard Start-Stop Assembly system [12] to generate a plasmid-encoded combinatorial metabolic pathway library for *C. acetobutylicum*. This would entail hierarchical library assembly in *E. coli*, using standard Start-Stop Assembly vectors for most steps, then for the final assembly step using a shuttle vector suitable both for *E. coli* and for transfer to the target organism, in this case *C. acetobutylicum*.

We began by generating sets of diverse promoters that would be needed. Libraries of synthetic promoters for *Clostridium* were previously generated [25] by introduction of 3-17 random mutations into the *C. acetobutylicum* thiolase gene promoter (P_*thl*_), resulting in synthetic promoters with a wide range of transcription strengths (260-fold range, from 0.7% to 178% activity of P_*thl*_). Anecdotal results from our lab and others indicate that *Clostridium* promoters are often very strong in *E. coli*, which can cause deleterious effects in *E. coli* cells, making DNA constructs containing such promoters difficult to construct and maintain in *E. coli*, and negatively affecting the diversity and quality of combinatorial libraries assembled in *E. coli*. Therefore, we tested whether fifteen promoters from two libraries previously characterised in *C. acetobutylicum* [25] pPM36-79% and pPM36-58% (where the % number is the chance of each position varied in the library matching the parent promoter in each variant) showed activity in *E. coli* by performing glucuronidase reporter assays (**Figure 1**). All tested promoters were active in *E. coli* and the set displayed a much narrower distribution of strengths (2-fold change, with the weakest and strongest promoters showing respectively 72% and 139% activity of P_*thl*_) than the same promoters in *C. acetobutylicum*.

**Figure 1.**
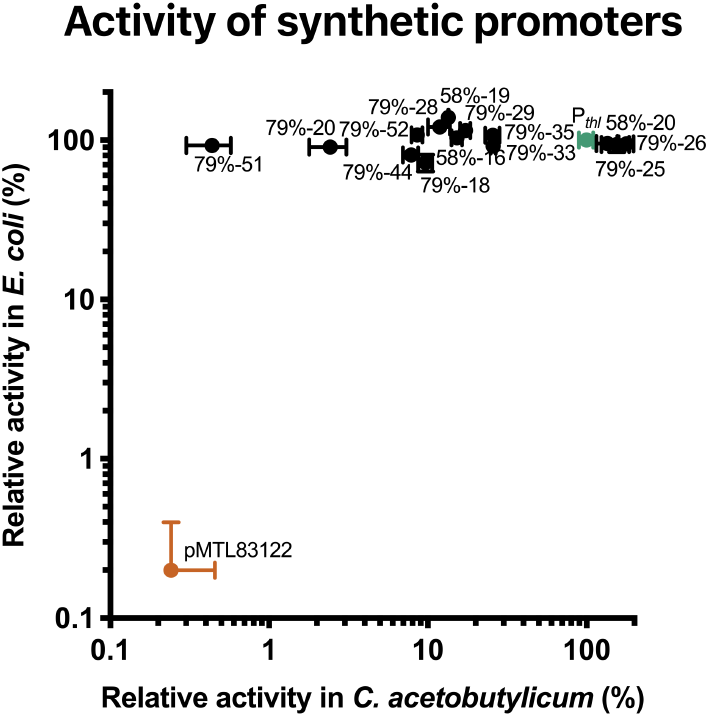
Activity of selected synthetic promoters from pPM36-79% and pPM36-58% plasmid libraries in *C. acetobutylicum* ATCC 824 and *E. coli* NEB5-alpha. Reporter glucuronidase activity of mid-exponential phase cells was normalized to the activity of a positive control strain transformed with plasmid pPM36-Pthl (*gusA* under P_*thl*_). A strain containing empty vector pMTL83122 was used as a negative control. Data is presented using a logarithmic scale. Error bars represent standard deviations of three independent experiments.

Next, we tested the feasibility of using *Clostridium* synthetic promoters for combinatorial assembly. Six synthetic promoters from the 79% library (79%-26, −25, −33, −29, 52 and −20) showing 70-fold change between the strongest and the weakest promoters were chosen and cloned in standard Start-Stop Assembly Level 0 format, meaning as α-β fusion site parts in the pStA0 storage plasmid (**Table S1**). These promoter parts were then used in Level 1 assembly reactions with previously-reported synthetic RBSs R1-R6 (plasmids pGT330-335), terminators (plasmids pGT337-340) [12] and a variety of different CDSs (*eyfp, adhE2, bktb* and *ter*; plasmids pGT431, pPM921, pPM924 and pPM934 respectively). After assembly reactions, the assembly reaction products were used to transform *E. coli* NEB5-alpha, and transformed cells were plated on LB agar supplemented with X-gal. Blue-white screening of the transformants showed that the apparent efficiency of Level 1 plasmid assembly was very low (0-2% of white colonies per plate), whereas non-expression controls in which the promoter parts were replaced with a short spacer sequence (Spacer 1 α-β [12]) showed much higher assembly efficiency (>98% white colonies) typical for Start-Stop Assembly [12]. This suggested that assembly was occurring normally, but constructs with burdensome strong synthetic *Clostridium* promoters were not viable in *E. coli*, so transformants containing those constructs did not grow to form white colonies.

The above observations indicate that *Clostridium* promoters which are too strong in *E. coli*, including the synthetic promoters studied here, are unsuitable for standard combinatorial assembly, because the complete expression units formed at intermediate steps (Levels 1, 2 and 3 in Start-Stop Assembly) in *E. coli* would be too burdensome, preventing the construction of satisfactory combinatorial libraries.

### A new approach: Integration-coupled activation of promoterless sequences (ICAPS)

As it appears that natural and synthetic promoters suitable for *Clostridium* often cause high burden in *E. coli*, this represents a general barrier to standard combinatorial assembly of libraries for *Clostridium*. Therefore, we set out to devise an alternative approach to overcome this barrier.

When using standard Start-Stop Assembly to construct operons, only the Level 1 expression unit for the first CDS in each operon requires a promoter, and only the Level 1 expression unit for the last CDS in each operon requires a terminator. Other (mid-operon) CDSs do not receive a promoter (or terminator) at Level 1 assembly. As a side effect, such mid-operon CDSs are not transcribed and therefore do not cause metabolic burden until a later assembly step (Level 2 or 3, depending upon the design), when they are placed downstream of a promoter. For our new approach, we generalised this effect by using a design in which every CDS, crucially even the first CDS, lacks a promoter or terminator, so the complete hierarchical assembly results in an entirely promoterless operon. This approach avoids transcription and thus associated metabolic burden throughout. Each CDS still receives an RBS, and mixtures of RBSs of varying strengths can be used to generate combinatorial libraries in which expression levels are varied. To be functional, these promoterless operon constructs and libraries must ultimately be transcribed. To achieve this, our new approach combines promoterless operon assembly with allele-coupled exchange (ACE) [8], a two-step allele exchange method for chromosomal integration of DNA. We used a version of ACE in which the second (final) homologous recombination event activates a promoterless antibiotic resistance marker gene (*ermB*) by placing it downstream of a promoter located on the chromosome (*thl*), ensuring cells only acquire resistance to the corresponding antibiotic (erythromycin) if the desired recombination has occurred. By constructing combinatorial libraries as promoterless operons via Start-Stop Assembly in an ACE vector, we avoid expression and burden throughout the entire assembly process. The entire operon is then activated simultaneously by ACE integration into the chromosome of the target organism, downstream of a promoter (**Figure 2**). We call this approach integration-coupled activation of promoterless sequences (ICAPS).

**Figure 2.**
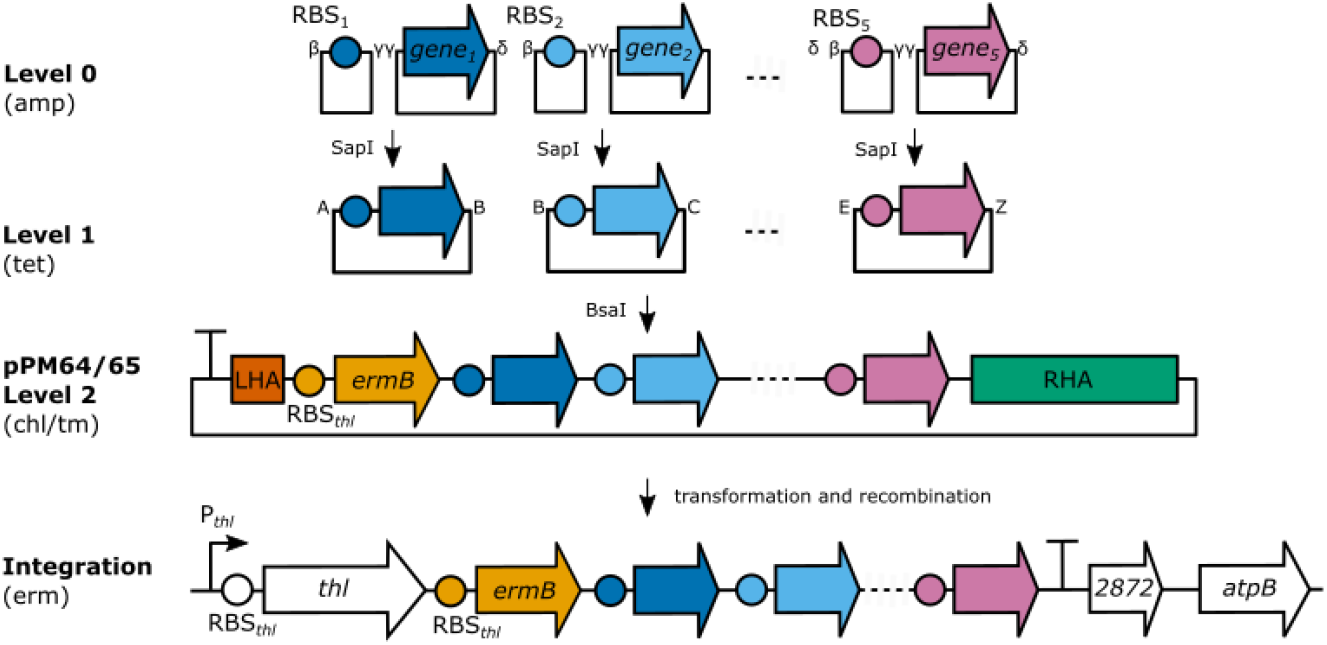
Integration-coupled activation of promoterless sequences (ICAPS) approach and ICAPS vector for *C. acetobutylicum*. RBS (β-γ) and CDS (γ-δ) parts are stored in standard Start-Stop Assembly Level 0 plasmids (resistant to ampicillin/carbenicillin). Level 1 plasmids (resistant to tetracycline) are generated by assembling RBS and CDS parts into modified Level 1 acceptor plasmids (β-δ, plasmids pPM902-pPM910). Up to five Level 1 plasmids and the new ICAPS Level 2 *E. coli-Clostridium* shuttle plasmids (pPM64, pPM65, chloramphenicol resistant) are used to generate Level 2 plasmids encoding promoterless operons encoding metabolic pathways. *C. acetobutylicum* is transformed with the Level 2 plasmid (selection with thiamphenicol). The pathway is activated by integration of the assembled promoterless operon into the bacterial chromosome downstream of a promoter (selection with erythromycin). See main text for details.

The existing Start-Stop Assembly system provides short spacer parts for use in place of promoters and/or terminators for construction of operons. Except for the special case of ICAPS in the present study, monocistronic designs are typically preferred, because varying both promoters and RBSs provides a wider range of expression levels, whereas operons provide options for less common cases such as linking the regulation of several CDSs using an operon design with a single regulated promoter. However, in ICAPS, every CDS will always lack a promoter and terminator, so if using standard Level 1 Start-Stop Assembly, spacers would be required instead of a promoter and terminator for every CDS. Therefore, to simplify, save effort and increase assembly efficiency, for ICAPS we constructed a special set of nine alternative Level 1 Start-Stop Assembly vectors omitting the standard promoter and terminator positions, thus with β-δ acceptor fusion sites instead of the standard α-ε acceptor fusion sites (pPM902-pPM910, **Figure 2, Table S1**). Level 1 assembly using these vectors generates Level 1 expression units without promoters or terminators, without requiring spacer parts, but is otherwise the same as standard Start-Stop Assembly, using standard Level 0 UTR/RBS (β-γ) and CDS (γ-δ) parts, and the same assembly procedure.

In order to implement ICAPS, combining promoterless operon assembly with ACE, we constructed special vector pPM64 (**Figure 2**), which contains a Level 2 Start-Stop Assembly cassette (the same as pStA212) with the relevant Level 2 (A-Z) acceptor fusion sites [12], in place of the previous multiple cloning site of *C. acetobutylicum* ACE vector pMTL-JH16 [8]. The key functional features (see Fig. 2) in 5’-3’ order are: the T1 terminator from CD0164 of *Clostridium difficile* 630 preventing transcriptional read-in from the backbone, a 300 bp fragment of the 3’ end of the *thl* gene serving as the left (second) homology arm (LHA) for chromosomal integration at the *C. acetobutylicum* ATCC 824 *thl* locus, a promoterless *ermB* gene conferring resistance to erythromycin only after integration, the Level 2 Start-Stop Assembly cassette from pStA212 and a 1200 bp right (first) homology arm (RHA).

Level 2 assembly is used to insert the promoterless and terminatorless Level 1 expression units into pPM64 in the designed order, resulting in a plasmid (or library) with a promoterless operon ready for integration and coupled activation. *C. acetobutylicum* is transformed with the assembled Level 2 plasmid or library by electroporation and transformants are selected on thiamphenicol. Initially, thiamphenicol-selected cells contain autonomous plasmids, but single-crossover integrants soon arise by homologous recombination at the large RHA, and grow more rapidly under thiamphenicol selection than the initial transformants which are limited by the plasmid’s pIM13 replicon [8]. Colonies are then re-streaked onto agar plates supplemented with erythromycin to select for cells in which a second recombination event at the short LHA has completed the double-crossover integration, placing the promoterless operon including *ermB* downstream of the strong chromosomal *thl* promoter, which is active in both acid and solvent production growth phases [8].

Initial experiments using pPM64 in Level 2 assembly showed frequencies of plasmid transformation into *C. acetobutylicum*, which were significantly lower than frequencies previously reported for other plasmids using the same pIM13 replicon, such as pMTL85151 [4]. Therefore, we tested if the low frequency could be a result of the RHA affecting the function of the Gram+ replication origin located downstream. We constructed pPM65 by inserting the TT2 terminator from the *fdx* gene of *C. pasteurianum* into pPM64 between the RHA and pIM13 replicon. The transformation frequency of the new plasmid pPM65 was improved 3 times when compared with pPM64 and was similar to pMTL85151 (**Figure S1**).

### Construction and testing of synthetic RBSs using a context-resistant design

Synthetic promoters are often used to vary expression levels in combinatorial designs, but ICAPS reserves transcriptional control to ensure assembled sequences are not expressed during assembly, and are activated upon integration, so promoters cannot be used to vary expression levels of CDSs in ICAPS. Instead, translational control can be used, which requires RBSs with a range of translation initiation rates.

Previously, a library of *Clostridium* synthetic parts allowing 20-fold change of the translation initiation rates was generated by changing the length of the spacer located between the Shine-Dalgarno (SD) sequence and the start codon in the reporter construct [26]. Another way of generating such libraries is randomisation of sequences flanking the consensus SD sequence, which has been successfully used in other organisms [12,27]. However, these strategies result in RBS parts with highly variable lengths or sequences, and the relative translation efficiency of these RBS parts may be context-dependent, meaning significantly affected by the CDS [28]. Therefore, we set out to generate a library of highly similar, ‘context-resistant’, synthetic RBSs showing a wider range of translation initiation ranges than the library reported before [26].

As a starting point for generation of synthetic RBSs, we used a 16 bp fragment of the expression plasmid pMTL83122 encoding the RBS of the *thl* gene with the NdeI restriction site scar aligning with the start codon (**Figure 3a**). This RBS was previously shown to allow significant expression of a number of genes [8,25,29]. In order to generate a set of RBSs with different translation initiation ranges we decided to randomise only two consecutive bases, −1 and +1 base of the Shine-Dalgarno AGGAGGT sequence, assuming that these changes will impact on the base-pairing interaction between the mRNA and the 16S rRNA and the translation initiation process, but the small number of differences will limit context-dependence of the RBS variants (**Figure 3a**). Synthetic RBSs were generated as Start-Stop Assembly Level 0 parts (with β-γ sites, standard for RBSs) by inverse PCR using empty pStA0 vector as a template and primers which introduce RBS sequences, giving a set of seven plasmids (pPM926-932) with different mutations in the RBS. These synthetic RBSs were assembled with the FLAG-*gusA* reporter CDS (from pPM911) in Level 1 assemblies, which were each used to transform *E. coli*, then plasmid DNA was prepared for each and used in Level 2 assembly with Level 2 ICAPS vector pPM64, resulting in the pPM71 plasmid library of promoterless RBS-FLAG-*gusA* reporter constructs ready for integration. Constructs were methylated *in vitro*, transformed into *C. acetobutylicum* and then integrated into the chromosome by selection on erythromycin and activation of the *ermB* marker (**Figure 3b**). Plasmid integrations were confirmed by PCR and sequencing. Glucuronidase activity assays of mid-exponential phase cultures of recombinant strains showed 65-fold change in the *gusA* expression between the strongest and the weakest RBS, O2 and O11 respectively (**Figure 3c**). The strongest RBS, O2, improved glucuronidase expression 3.7 times when compared with the Start-Stop Assembly-formatted RBS from pMTL83122 plasmid (O20). Therefore, we successfully generated a library of highly-similar RBS parts (which are likely context-resistant) showing a wide range of translation initiation ranges in *C. acetobutylicum* tested using a reporter system.

**Figure 3.**
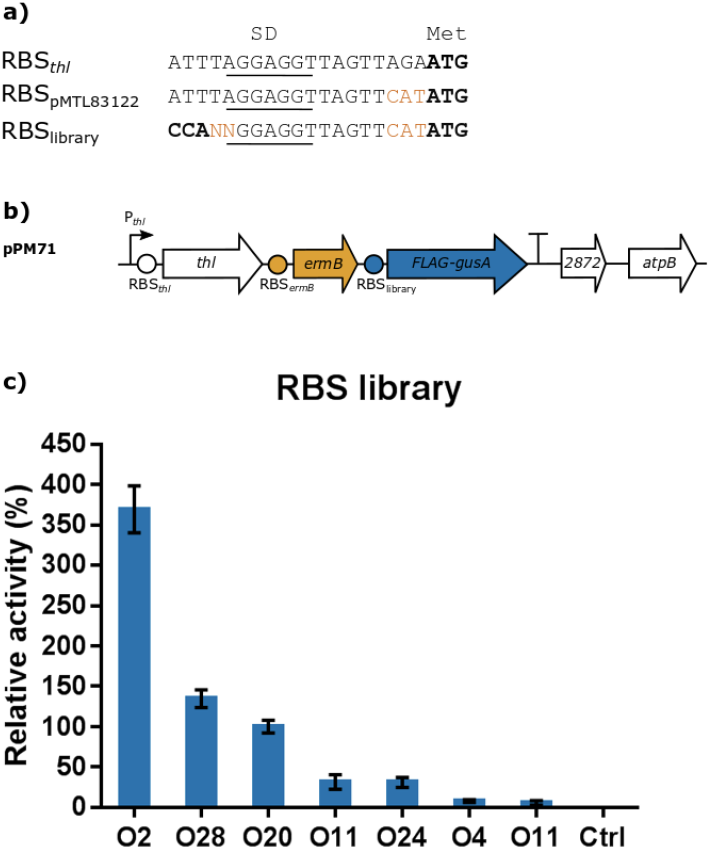
Context-resistant synthetic RBS library in *C. acetobutylicum*. (**a**) Nucleotide sequences of RBSs in the *thl* gene, the pMTL83122 expression plasmid and the synthetic RBS library. SD sequence is underlined. Start codon (ATG, Level 1 β-fusion site) and Level 1 γ-fusion site (CCA) are marked in bold. CAT resulting from the introduction of the NdeI restriction site is marked in orange. The context-resistant synthetic RBS library was generated by randomisation of −1 and +1 bases of SD (NN marked in orange). (**b**) Locus of the *thl* gene after integration of the pPM71 plasmid library encoding context resistant synthetic RBSs and FLAG-*gusA* reporter gene. (**c**) Relative glucuronidase activity of mid-exponential phase cultures with integrated context resistant synthetic RBSs (O1, O2, O4, O11, O20, O28) and FLAG-*gusA* reporter gene. Activities were normalised to the activity of the strain encoding RBS O20 (Start-Stop adapted RBS from pMTL83122 plasmid). Strain transformed with the empty pPM64 integration plasmid was used as a negative control (Ctrl). Error bars represent standard deviations of three independent experiments.

**Figure 4.**
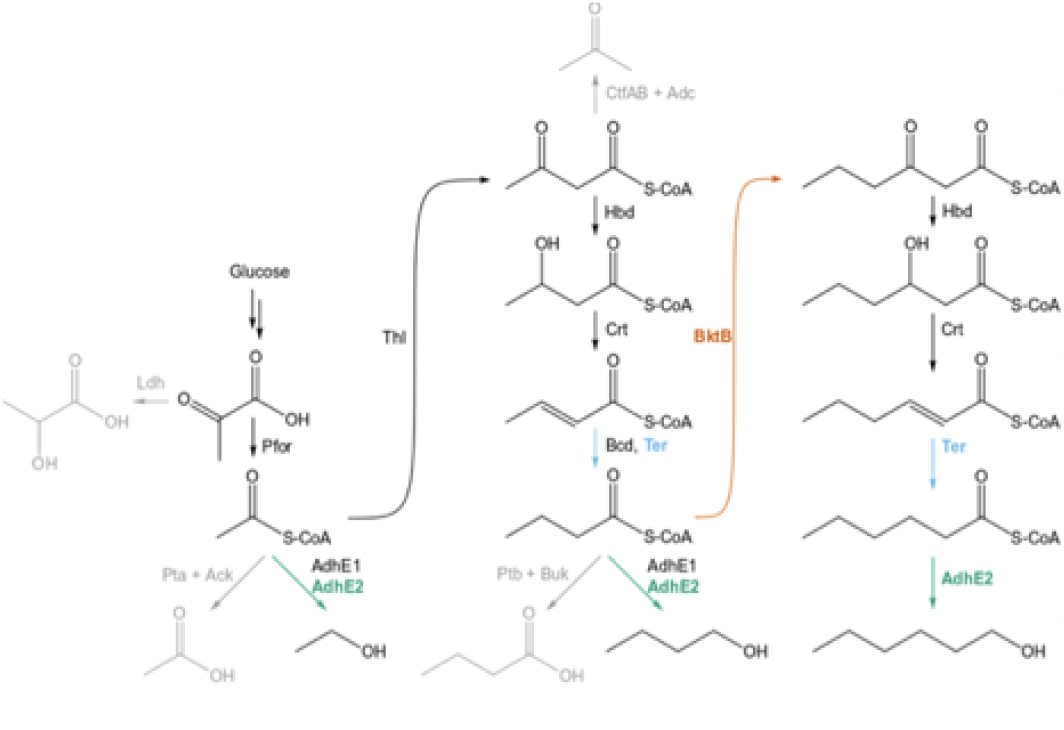
Recombinant 1-hexanol pathway in *C. acetobutylicum*. The native butanol pathway is extended to produce a six carbon chain alcohol by recombinant expression of β-ketothiolase (BktB) from *C. necator* H16, *trans*-enoyl CoA reductase from *Treponema denticola* (Ter) and overexpression of *C. acetobutylicum* bifunctional aldehyde-alcohol dehydrogenase (AdhE2). Other enzymes involved are annotated as follows: lactate dehydrogenase (Ldh), pyruvate:ferredoxin oxidoreductase (Pfor), phosphotransacetylase (Pta), acetate kinase (Ack), bifunctional aldehyde-alcohol dehydrogenase (AdhE1), acetoacetate decarboxylase (Adc), coenzyme A transferase (CtfAB), 3-hydroxybutyryl-CoA dehydrogenase (Hbd), short-chain-enoyl-CoA-hydratase (Crt), butyryl-CoA dehydrogenase (Bcd), phosphotransbutyryalse (Ptb), butyrate kinase (Buk).

We also tested the strengths and the range of synthetic RBSs in the *E. coli* expression system. Plasmid library pPM71 contained promoterless synthetic RBS-FLAG*-gusA* expression units which could therefore not be used to determine translation initiation rates in *E. coli*. An additional set of Level 1 plasmids (plasmid library pPM87) was assembled in which the synthetic RBS-*eyfp* expression unit was transcribed from the inducible *rhaBAD* promoter. All seven synthetic RBSs were strong when compared with the *E. coli* RBSs R1-R6 characterised previously [12] (**Figure S2**). The set of RBSs showed a much narrower range of strengths in *E. coli* than in *C. acetobutylicum*, only 1.7 fold change between the strongest and the weakest RBSs (O28 and O1, respectively).

### Assembly and integration-coupled activation of promoterless hexanol pathway operon

To test the ICAPS approach and the new RBSs, we attempted to engineer the CoA-dependant alcohol pathway in *C. acetobutylicum*, to extend it to produce 1-hexanol. Previously, production of 1-butanol in *E. coli* was achieved by overexpression of *C. acetobutylicum* CoA-dependent pathway genes (*thl, hbd, crt, bcd-etfAB* and *adhE2*) [30,31] and further improved by creating NADH and acetyl-CoA driving forces and replacing the acyl-CoA dehydrogenase (Bcd) with NADH-dependent *trans*-enoyl-CoA reductase (Ter) from *Treponema denticola*[32]. Subsequently, the 1-butanol pathway was extended to 1-hexanol, a six carbon chain product, by overexpression of the β-ketothiolase (Bktb) from *Cupriavidus necator* H16 showing specificity for condensation of longer carbon chain substrates[19].

The CDSs of *bktb* and *ter* were codon-optimised for expression in *C. acetobutylicum*, and the CDS of *adhE2* was recoded (the sequence was similarly regenerated with suitable codon usage) to avoid undesired recombination between the introduced copy of the CDS and the native *adhE2* gene on the *C. acetobutylicum* native pSOL1 plasmid. All three CDSs were synthesised as linear DNA fragments (gBlocks, Integrated DNA Technologies) and inserted into Start-Stop Assembly Level 0 plasmid resulting in plasmids pPM921, pPM924 and pPM934. CDSs were combinatorially assembled with six synthetic RBSs (O2, O4, O11, O20, O24 and O28) in Level 1 plasmids pPM902 (AB), pPM904 (BC) and pPM907 (CZ) and used to transform *E. coli* to generate Level 1 plasmid libraries. White colonies from transformation plates were pooled and plasmid DNA from each Level 1 library was purified. Randomisation of the RBS sequences was confirmed by DNA sequencing (**Figure S3a**). Level 1 plasmid libraries were then assembled together in the Level 2 ICAPS vector pPM65 to generate Level 2 plasmid library pPM76, encoding the full-length promoterless combinatorial 1-hexanol pathway. The fidelity of the pPM76 assembly was assessed by picking 10 random white colonies from the transformation plate and restriction analysis of the purified plasmid DNA (**Figure S3b**). All tested plasmids showed the expected band pattern showing that the pPM76 plasmid library contains correctly-assembled expression units. The remaining white colonies from pPM76 were pooled and used for plasmid DNA purification. The plasmid DNA was methylated and used to transform *C. acetobutylicum*. The empty pPM65 vector was also transformed to generate control strains not containing the recombinant pathway. Transformants were selected by incubation on agar plates containing thiamphenicol, then colonies were re-streaked onto agar plates containing erythromycin in order to select for integrants in which integration-coupled activation of *ermB* and the assembled promoterless hexanol operon had occurred (**Figure 5a**). Erythromycin-resistant colonies were used to inoculate 10 mL liquid cultures in CBMS medium supplemented with erythromycin and grown for 72 h. The concentration of ethanol, butanol and 1-hexanol was quantified by GC-MS (**Figure S4**). 33 out of 40 strains showed visible growth and alcohol production. Six strains (76-16, −17, −18, −20, −25 and −31) produced detectable amounts of 1-hexanol (from 0.026 mM to 0.048 mM). Interestingly, two other strains (76-2 and −6) showed increased titers of ethanol when compared to the control strains (65-1, −2 and −3). Genomic DNA from selected strains was purified and used as a template for PCR to identify RBSs for each coding sequence in the combinatorial pathway (**Figure 5b**). Genotyping showed that the top three 1-hexanol producers had the strongest synthetic RBS O2 in the *bktb* expression unit, whereas distribution of synthetic RBSs for other genes was more diverse. These results may suggest that condensation of acetyl-CoA and butyryl-CoA to 3-hydroxyhexanoyl-CoA is the rate limiting step in the 1-hexanol pathway. Analysis of RBSs in the ethanol hyper-producing strains showed that in strain 76-2 all CDSs were expressed from weak RBSs (O4 and O24), whereas strain 76-6 had strong RBSs for all three CDSs (O2 and O28). However, the *bktb* coding sequence in this strain 76-6 was interrupted with a 2 kb fragment of the plasmid backbone, which could explain lack of 1-hexanol production.

**Figure 5.**
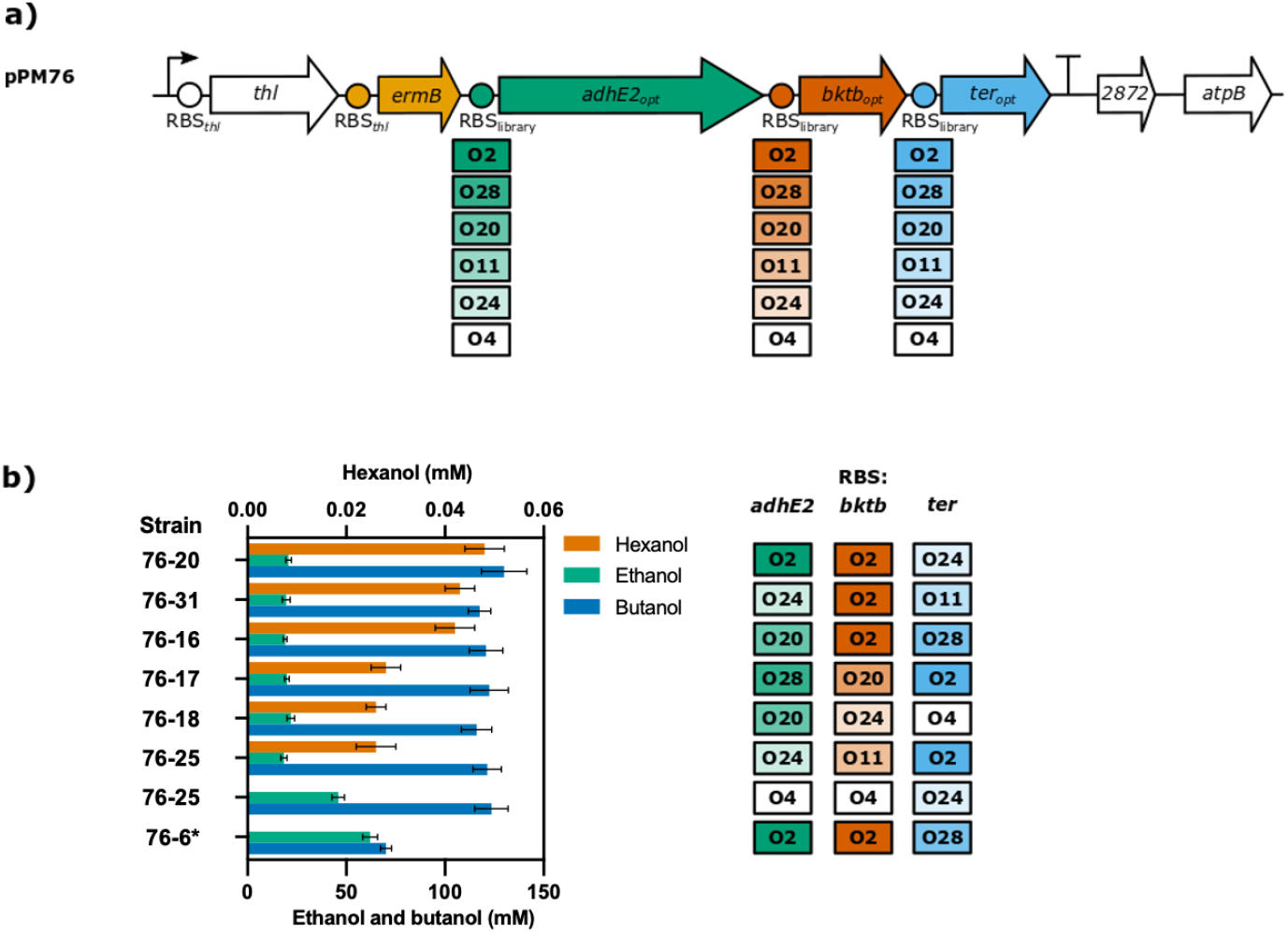
Recombinant 1-hexanol pathway in *C. acetobutylicum*. **(a)** Locus of the *thl* gene after integration of the pPM76 plasmid library encoding the combinatorially assembled pathway consisting of *adhE2, bktb* and *ter* genes. Six RBSs of different strengths (O2, O4, O11, O20, O24 and O28) were used for expression of each gene. The intensity of the colour represents the strength of the RBS (based on the assay with the FLAG-*gusA* reporter) **(b)** Fermentation experiment using the recombinant strains. Forty erythromycin-resistant strains were grown in CBMS for 72 h. Integrants of the empty pPM65 plasmid were used as controls. Concentrations of ethanol, butanol and hexanol in the culture medium were quantified using GC-MS. Results are presented for strains producing detectable amounts of hexanol or showing increased production of ethanol or butanol. RBSs for each gene were determined by PCR and DNA sequencing. Strain marked with * (76-6) showed unexpected length of the *bktb* PCR product and large rearrangements in the coding sequence.

## Discussion

Combinatorial design and construction using multi-part DNA assembly is a very useful approach to quickly and easily obtain effective designs of DNA constructs encoding metabolic pathways or other systems. However, in standard designs, hierarchical DNA assembly systems generate intermediate constructs with complete expression units, leading to expression of CDSs and metabolic burden in the typical assembly host *E. coli*, reducing library size, quality, and exploration of design space. Here we tested *Clostridium* synthetic promoters in *E. coli*, and found they were too strong and thus burdensome in *E. coli* to allow standard combinatorial assembly, in keeping with previous anecdotal observations. This issue may be a general barrier to combinatorial assembly for *Clostridium* and other organisms. To address this problem, here we developed integration-coupled activation of promoterless sequences (ICAPS), assembling constructs or libraries as promoterless operons which are inactive during assembly, and activating their expression only upon ACE chromosomal integration downstream of a promoter.

As an ICAPS proof of concept, we constructed a combinatorial library for an extended CoA-dependent alcohol pathway in *C. acetobutylicum* and activated it via ACE integration, achieving detectable hexanol production in a strain not previously engineered for this product. Functional variants were obtained at a good frequency (15% of colonies screened) from a relatively small library of 216 possible designs, indicating the usefulness of the approach. Our results also identified *bktb* as a potential bottleneck, showing how combinatorial approaches can readily reveal expression trends and targets for improvement.

The titres we observed were modest compared to native producers such as *C. carboxidivorans* [21,22] or engineered *C. ljungdahlii* [24]. This suggests that improvements could be made, and that the range of expression levels used did not maximise production of hexanol. The system could be improved by developing further RBSs with higher expression levels, and by testing further chromosomal integration sites with promoters providing different levels and patterns of expression.

ICAPS, and ACE integration which ICAPS builds upon, have similarities with classic promoter trap/gene trap methods, which use transcriptional activation of a reporter or marker gene upon chromosomal integration of a recombinant transposon to identify active promoters [33]; and the more recent Tn-Seq, a high-throughput parallel method enabling fitness and genetic interaction studies [34] and identification of essential genes [35]. Here we applied a similar principle but in a synthetic biology context, to facilitate design and construction, rather than analysis.

New vectors and parts constructed for this study include a special set of nine alternative Start-Stop Assembly Level 1 vectors for simple, efficient assembly of promoterless operons (omitting the standard promoter and terminator positions, thus with β-δ acceptor fusion sites); a set of context-resistant synthetic RBSs, and *E. coli*–*C. acetobutylicum* ICAPS shuttle vector pPM65. These vectors and parts may be useful to others either directly, or as an example of how the ICAPS approach could be implemented in other cases.

## Materials and methods

### Bacterial strains and growth conditions

*E. coli* NEB5-alpha (NEB) was used for plasmid construction and was grown in lysogeny broth (LB) at 37°C with rotary shaking at 250 rpm or on LB agar plates. *E. coli* strains transformed with plasmids (Table S1) were cultured in LB broth or on LB plates supplemented with carbenicillin (100 μg·mL^-1^), tetracycline (10 μg·mL^-1^) or chloramphenicol (12.5 μg·mL^-1^). *E. coli* blue/white colony screening was performed on LB agar plates supplemented with ChromoMax™ IPTG/X-Gal Solution (Fisher BioReagents). *Clostridium acetobutylicum* ATCC 824 was maintained at 37°C under an anaerobic atmosphere of N_2_:H_2_:CO_2_ (80:10:10, vol:vol:vol) in a Whitley A35 Anaerobic Workstation (Don Whitley, UK) in *Clostridium* Basal Medium (CBM) static cultures or on CBM agar plates [36]. Before inoculation, media were prereduced overnight in the anaerobic workstation. Cultures were supplemented with thiamphenicol (15 μg·mL^-1^) as required for plasmid selection and with erythromycin (40 μg·mL^-1^) for selection of genomic integrants.

### Plasmid construction and combinatorial assembly

Plasmid construction was carried out using standard molecular biology methods [37]. Construction of the new Start-Stop Assembly plasmids (pPM900-pPM910, pPM64-lacZ and pPM65-lacZ) and Level 0 plasmids encoding the synthetic RBS library (plasmids pPM926-pPM932), FLAG*-gusA* (pPM911), *adhE2* (pPM921), *bktb* (pPM924) and *ter* (pPM934) is described in the Supplementary Materials.

Plasmid libraries pPM71 and pPM76 were assembled by assembly reactions containing 20 fmol of destination plasmid, 60 fmol of each insert, T4 DNA Ligase (400 U; NEB), T4 DNA Ligase buffer (1x; NEB) and the appropriate restriction enzyme (SapI or BsaI, 10 U; NEB). Reactions were incubated in a thermocycler for 30 two-step cycles of 37°C for 5 min then 16°C for 5 min, before a final digestion step at 42°C for 5 min and a denaturation step at 65°C for 20 min.

Plasmid library pPM71 (pPM71-O1, O2, -O4, -O11, -O20, -O24 and -O28) was generated by performing individual assembly reactions for each plasmid. Each Level 1 reaction contained the appropriate RBS part (pPM926-pPM932), FLAG*-gusA* donor plasmid (pPM911) and the recipient plasmid pPM903 (1AZ, β-δ). Reaction mixtures were used to transform *E. coli*. Level 1 plasmids purified from cultures originating from single colonies from the transformation plates were used in individual Level 2 reactions with the recipient plasmid pPM64-lacZ.

Plasmid library pPM76 was generated by a combinatorial assembly of RBS parts for each coding sequence. Level 1 reaction AB contained an equimolar mixture of six RBS donor plasmids (pPM927-pPM932), *adhE2* CDS (pPM921) and the recipient plasmid pPM902 (1AB, β-δ). Reaction BC contained an equimolar mixture of six RBS donor plasmids (pPM927-pPM932), *bktb* CDS (pPM924) and the recipient plasmid pPM904 (1BC, β-δ). Reaction CZ contained an equimolar mixture of six RBS donor plasmids (pPM927-pPM932), *ter* CDS (pPM934) and the recipient plasmid pPM907 (1CZ, β-δ). Reaction mixtures were used to transform *E. coli* and plated on selection plates supplemented with X-Gal and IPTG. White colonies from each transformation plate were collected using an inoculation loop and pooled together for plasmid purification to generate three Level 1 plasmid libraries. Then, the three Level 1 plasmid libraries were used in a single assembly reaction with the recipient plasmid pPM65-lacZ and the reaction mixture was transformed to *E. coli*. White colonies from the transformation plate were collected using an inoculation loop and pooled together for plasmid purification.

### Clostridium acetobutylicum transformation and chromosome integration

Plasmids or plasmid libraries (3 μg) were methylated *in vitro* using GpC methyltransferase M.CviPI (NEB), transformed to *C. acetobutylicum* by electroporation [4] and plated on 2YTG agar plates (tryptone, 16 g/L; yeast extract, 10 g/L; NaCl, 5 g/L; glucose, 20 g/L; pH 5.2) supplemented with thiamphenicol (15 μg·mL^-1^). Single colonies from transformation plates were streaked on CBM agar plates supplemented with erythromycin (40 μg·mL^-1^) to select for genomic integrants. Erythromycin-resistant colonies were re-streaked on the same medium to confirm the phenotype.

### Glucuronidase assay

Glucuronidase activity in *E. coli* and *C. acetobutylicum* was determined as described by Dupuy and Sonenshein [38]. Briefly, strains expressing *gusA* were grown in CBM with erythromycin to OD 600 nm of 1 (Eppendorf BioSpectrometer kinetic). Samples (1.5 mL) were harvested by centrifugation and pellets were frozen at −80°C. Before testing, pellets were resuspended in 0.8 mL of buffer Z (60 mM Na_2_HPO_4_·7H_2_O, 40 mM NaH_2_PO_4_·H_2_O, 10 mM KCl, 1 mM MgSO_4_·7H_2_O, pH adjusted to 7.0, and 50 mM 2-mercaptoethanol added freshly). 0.2 mL of each sample was used for OD 600 nm measurement. To the remaining sample (0.6 mL), toluene (6 μL) was added, tubes were vortexed for 1 min and incubated on ice for 10 min. Tubes were transferred into a 37 °C heating block and preincubated for 30 min with caps open. Reactions were started by addition of 120 μL of 6 mM *p*-nitrophenyl-β-D-glucuronide (Merck Millipore) solution in buffer Z. After incubation at 37°C (5-30 min), reactions were stopped by addition of 1 M Na_2_CO_3_ (300 μL) and reaction time was recorded. Cell debris was removed by centrifugation at 10000 x g for 10 min. Supernatants were transferred into polystyrene spectrophotometer cuvettes and absorbance at 405 nm was determined (Eppendorf BioSpectrometer kinetic). Relative glucuronidase activity was calculated by dividing the absorbance at 405 nm by sample OD 600 nm and incubation time and normalization to the positive control sample (O20, RBS_*thl*_).

### Extended carbon chain alcohol fermentation and strain genotyping

Cultures in CBMS [39] (10 mL) supplemented with erythromycin (40 μg·mL^-1^) were inoculated with fresh colonies from integration plates and incubated at 37°C for 72 h. Samples (1 mL) were collected and cells were separated by centrifugation at 10,000 x *g* for 3 min. Supernatants (0.5 mL) were filtered through a Nylon syringe filter (0.22 μm, 13 mm; Thames Restek UK) and extracted with ethyl acetate (0.5 mL) containing an internal standard *tert*-butylbenzene (10 mM). Ethanol, butanol, hexanol and octanol were quantified using a gas chromatograph (Agilent 7890B) equipped with a DB-624 Ultra Inert capillary column (30 m by 0.25 mm by 1.4 μm; Agilent) and a mass selective detector (Agilent 5977A). Ultrapure helium was used as carrier gas at 0.8 mL min^-1^ flow rate. Samples (0.2 μL, split ratio 100:1) were injected at 240 °C. The initial oven temperature was held at 35°C for 6 min, then increased at 10 °C min^−1^ to 260 °C, and held for 1 min. Fermentation products were quantified by comparing their peak areas normalised to the peak area of the internal standard with calibration curves of authentic standards (Sigma). Cell pellets separated before ethyl acetate extraction were used for purification of genomic DNA [40]. Extracted DNA (50 ng) was used for PCR with primers oligoPM326 and oligoPM455 (amplification of the left integration junction, including the RBS of *adhE2* gene), oligoPM446 and oligoPM327 (3’ end of *adhE2, bktb* and 5’ end of *ter*, including RBSs of *bktb* and *ter* genes) and oligoPM456 and oligoPM324 (right integration junction). Gel-purified PCR products were sequenced with primers oligoPM550 (RBS_*adhE2*_), oligoPM446 (RBS_*bktb*_) and oligoPM447 (RBS_*ter*_).

## Acknowledgments

The authors thank Dr Soo Mei Chee at Imperial for GC-MS analysis, and Dr Alexandra Faulds-Pain for critical reading of the manuscript.

## Author contributions

PM and JH designed the study. PM performed the experiments. JH supervised the work. PM, JH and JW all wrote and revised the manuscript.

## Data availability

The supplementary information includes details of plasmid construction, Table S1 Plasmids used in the study, Table S2 Oligonucleotides used in the study, and supplementary figures.

## Material availability

Materials are available from the authors upon reasonable request.

