## Supplementary Information for "Integration-coupled activation of promoterless combinatorial pathway libraries in *Clostridium* avoids burden during DNA assembly"

**Table S1.** Plasmids used in the study.

| Plasmid | StA level | Details | Marker | Accession Number | Reference |
|---|---|---|---|---|---|
| **Empty Start-Stop Assembly (StA) plasmids** | | | | | |
| pStA0 | 0 (*lacZ*) | | *ampR* | MG649420 | [1] |
| pPM900-ccdB | 0 (*ccdB*) | | *ampR* | MT361981 | This study |
| pStA1AZ | 1 AZ (α-ε) | | *tetR* | MG649422 | [1] |
| pStA1BZ | 1 BZ (α-ε) | | *tetR* | MG649424 | [1] |
| pStA1CZ | 1 CZ (α-ε) | | *tetR* | MG649426 | [1] |
| pStA1DZ | 1 DZ (α-ε) | | *tetR* | MG649428 | [1] |
| pStA1EZ | 1 EZ (α-ε) | | *tetR* | MG649429 | [1] |
| pPM901 | 1 AB (α-δ) | | *tetR* | MT361982 | This study |
| pPM902 | 1 AB (β-δ) | | *tetR* | MT361983 | This study |
| pPM903 | 1 AZ (β-δ) | | *tetR* | MT361984 | This study |
| pPM904 | 1 BC (β-δ) | | *tetR* | MT361985 | This study |
| pPM905 | 1 BZ (β-δ) | | *tetR* | MT361986 | This study |
| pPM906 | 1 CD (β-δ) | | *tetR* | MT361987 | This study |
| pPM907 | 1 CZ (β-δ) | | *tetR* | MT361988 | This study |
| pPM908 | 1 DE (β-δ) | | *tetR* | MT361989 | This study |
| pPM909 | 1 DZ (β-δ) | | *tetR* | MT361990 | This study |
| pPM910 | 1 EZ (β-δ) | | *tetR* | MT361991 | This study |
| pPM64-lacZ | 2 | | *catP* | MT361992 | This study |
| pPM65-lacZ | 2 | | *catP* | MT361993 | This study |
| **Level 0 plasmids** | | | | | |
| pGT330 | 0 (β-γ) | RBS R3 (c13) | *ampR* | MG649441 | [1] |
| pGT331 | 0 (β-γ) | RBS R2 (c33) | *ampR* | MG649442 | [1] |
| pGT332 | 0 (β-γ) | RBS R1 (c44) | *ampR* | MG649445 | [1] |
| pGT333 | 0 (β-γ) | RBS R4 (c58) | *ampR* | MG649446 | [1] |
| pGT334 | 0 (β-γ) | RBS R5 (c36) | *ampR* | MG649443 | [1] |
| pGT335 | 0 (β-γ) | RBS R6 (c42) | *ampR* | MG649444 | [1] |
| pGT337 | 0 (δ-ε) | T1 (L3S2P55) | *ampR* | MG649450 | [1] |
| pGT338 | 0 (δ-ε) | T2 (L3S2P21) | *ampR* | MG649449 | [1] |
| pGT339 | 0 (δ-ε) | T3 (ECK120033737 | *ampR* | MG649448 | [1] |
| pGT340 | 0 (δ-ε) | T4 (ECK120019600) | *ampR* | MG649447 | [1] |
| pGT431 | 0 (γ-δ) | pStA0::*eyfp* | *ampR* | - | [1] |
| pPM911 | 0 (γ-δ) | FLAG-*gusA* | *ampR* | MT361994 | This study |
| pPM921 | 0 (γ-δ) | *adhE2*<br>*C. acetobutylicum*<br>codon optimised | *ampR* | MT361995 | This study |
| pPM924 | 0 (γ-δ) | *bktb*<br>*C. necator* H16<br>codon optimised | *ampR* | MT361996 | This study |
| pStA0-RBS | 0 (α-γ) | $RBS_{thl}$ | *ampR* | - | This study |
| pPM926 | 0 (β-γ) | RBS O1 | *ampR* | MT361997 | This study |
| pPM927 | 0 (β-γ) | RBS O2 | *ampR* | MT361998 | This study |
| pPM928 | 0 (β-γ) | RBS O4 | *ampR* | MT361999 | This study |
| pPM929 | 0 (β-γ) | RBS O11 | *ampR* | MT362000 | This study |
| pPM930 | 0 (β-γ) | RBS O20 (thl) | *ampR* | MT362001 | This study |
| pPM931 | 0 (β-γ) | RBS O24 | *ampR* | MT362002 | This study |
| pPM932 | 0 (β-γ) | RBS O28 | *ampR* | MT362003 | This study |
| pPM934 | 0 (γ-δ) | *ter*<br>*Treponema denticola*<br>codon optimised | *ampR* | MT362004 | This study |
| pPM957 | 0 (α-β) | $P_{rhaBAD}$ | *ampR* | MT362005 | This study |

| | | | | | |
|---|---|---|---|---|---|
| **Other plasmids** | | | | | |
| pMTL-JH16 | | ACE plasmid, pIM13 Gram+ origin | *catP* | HQ875757 | [2] |
| pCK302 | | | *ampR* | KU555410 | [3] |
| pUC19 | | | *ampR* | - | NEB |
| pPM71 library | 2 | AZ: synthetic RBS, FLAG-*gusA* | *catP* | - | This study |
| pPM76 library | 2 | AB: synthetic RBS library, *adhE2*<br>BC: synthetic RBS library, *bktb*<br>CZ: synthetic RBS library, *ter* | *catP* | - | This study |
| pPM87 library | 1 | AB: $P_{rhaBAD}$, synthetic RBS, *eyfp* | *tetR* | - | This study |

**Table S2.** Oligonucleotides used in the study. 5'-phosphorylated primers are denoted by /5Phos/.

| Name | Sequence | Description |
|---|---|---|
| oligoPM383 | GATAGCGGCCGCGGAGAGAGACCGGGCAGTGAGCGCAACGC | Construction of pPM64-lacZ |
| oligoPM384 | GATAGCTAGCAGTATGAGACCCTATGCGGCATCAGAGCAGATTG | |
| oligoPM434 | /5Phos/TTATGAAGAGCGGAAACTATG | Construction of pPM901-pPM910 |
| oligoPM435 | AATGAGAGACCATTTGTCCTACTC | |
| oligoPM436 | AGGTAGAGACCATTTGTCCTACTC | |
| oligoPM437 | GCTTAGAGACCATTTGTCCTACTC | |
| oligoPM438 | CGCTAGAGACCATTTGTCCTACTC | |
| oligoPM439 | TACTAGAGACCATTTGTCCTACTC | |
| oligoPM479 | CTAACCTCCTACTGCGAAGAGCCACACTGGATTCTCACCAATAAAAAACGCCCG | Construction of pStA0-RBS |
| oligoPM480 | TTCATATGTGAAGAGCGTAAGACCTCTAGGGCGGCGGATTTG | |
| oligoPM487 | AAGGAGGTTAGTTCATATGTGAAG | Construction of the RBS library in level 0 |
| oligoPM488 | AGGGAGGTTAGTTCATATGTGAAG | |
| oligoPM489 | TTGGAGGTTAGTTCATATGTGAAG | |
| oligoPM490 | CTGGAGGTTAGTTCATATGTGAAG | |
| oligoPM491 | GCGGAGGTTAGTTCATATGTGAAG | |
| oligoPM492 | CCGGAGGTTAGTTCATATGTGAAG | |
| oligoPM517 | TAGGAGGTTAGTTCATATGTGAAG | |
| oligoPM518 | /5Phos/TGGCGAAGAGCCACACTGG | |
| oligoPM506 | GTGAGAATCCAGTGTGAGAGACCGGCTTACTAAAAGCCAG | Amplification of the *ccdB* cassette |
| oligoPM507 | CGATGCTCTAGAGGTCTGAGACCTTATATTCCCCAGAACATCAG | |
| oligoPM519 | /5Phos/CCAAGAAGAGCCAGTAGGGCAG | Construction of pPM901-pPM910 |
| oligoPM520 | CTCCTGAGACCATTCTCACCAATAAAAAAC | |
| oligoPM521 | CATTTGAGACCATTCTCACCAATAAAAAAC | |
| oligoPM522 | ACCTTGAGACCATTCTCACCAATAAAAAAC | |
| oligoPM523 | AAGCTGAGACCATTCTCACCAATAAAAAAC | |
| oligoPM524 | AGCGTGAGACCATTCTCACCAATAAAAAAC | |
| oligoPM563 | TTGCAGGCTTCTTATTTTTATGGCGCGCCGCATTCACTTCTTTTC | Construction of pPM65-lacZ |
| oligoPM564 | ATGCAGGCTTCTTATTTTTATTTCTTTTTATTGCAGTTGCATTTATTAAAAATGCACTTACTAAAGCAAAG | |
| oligoPM617 | AAGGGGTTGGTCTCATGTGGCTCTTCGCAGCCACAATTCAGCAAATTGTGAAC | Construction of $P_{rhaBAD}$ |
| oligoPM618 | CAGTGTTGGGTCTCTGGTCGCTCTTCATGGAAGTTAAACAAAATTATTTCTAGAGGGAAAC | |
| oligoPM326 | gcttggagtaaaaccacttgc | Genotyping of pPM76 integrants |
| oligoPM455 | AGCTCATTTAGTTTCTGCTTTA | |
| oligoPM446 | GGGTACTTCAGACACTGAAAAGG | |
| oligoPM327 | TTGAAGCAAGTCCGTATCCA | |
| oligoPM456 | AGAAGGAATAAATTATGAAGCAGA | |
| oligoPM324 | ccatgaagaggtactggcaat | |
| oligoPM550 | tcaaaatgggtcaatcgaga | |
| oligoPM447 | AATGATGGAAAGGGCTGAAG | |
| oligoGT538 | CAGTGGTCAGCGACT | Promoter spacer |
| oligoGT539 | TGGAGTCGCTGACCA | |
| oligoPM326 | gcttggagtaaaaccacttgc | pPM71 integration screen and sequencing |
| oligoPM177 | CCAACGCTGATCAATTCCAC | |

### 1. Plasmid construction

#### 1.1. Generation of new empty StA plasmids

**pPM900-ccdB**
The 637 bp *ccdB* cassette from plasmid pDONR™/Zeo with the BsaI restriction site silenced (GGTCTC→GGTGTC) was custom synthesised as a gBlock Gene Fragment (IDT). The cassette was amplified with primers oligoPM506 and oligoPM507, digested with BsaI and ligated to pStA0 vector cut with the same enzyme. The ligation reaction mixture was used to transform One Shot™ *ccdB* Survival™ 2 $T1^R$ *E. coli* strain (Thermo Fisher).

**pPM901-pPM910**
Plasmid pPM901 was generated by inverse PCR (Q5 High-Fidelity DNA Polymerase, NEB) with primers oligoPM434 and oligoPM435 using pStA1AZ as template, DpnI (NEB) treatment and PCR product circularisation with T4 Ligase (NEB). Plasmid pPM902 was generated by inverse PCR with primers oligoPM519 and oligoPM520 using pPM901 as template, DpnI treatment and PCR product circularisation with T4 Ligase. Plasmid pPM903 was generated in two steps, by inverse PCR with primers oligoPM434 and oligoPM439 using pStA1AZ as template, DpnI treatment and PCR product circularisation with T4 Ligase, followed by inverse PCR with primers oligoPM519 and oligoPM520, DpnI treatment and PCR product circularisation with T4 Ligase. Plasmid pPM904 was generated in two steps, by inverse PCR with primers oligoPM434 and oligoPM436 using pStA1BZ as template, DpnI treatment and PCR product circularisation with T4 Ligase, followed by inverse PCR with primers oligoPM519 and oligoPM521, DpnI treatment and PCR product circularisation with T4 Ligase. Plasmid pPM905 was generated in two steps, by inverse PCR with primers oligoPM434 and oligoPM439 using pStA1BZ as template, DpnI treatment and PCR product circularisation with T4 Ligase, followed by inverse PCR with primers oligoPM519 and oligoPM521, DpnI treatment and PCR product circularisation with T4 Ligase. Plasmid pPM906 was generated in two steps, by inverse PCR with primers oligoPM434 and oligoPM437 using pStA1CZ as template, DpnI treatment and PCR product circularisation with T4 Ligase, followed by inverse PCR with primers oligoPM519 and oligoPM522, DpnI treatment and PCR product circularisation with T4 Ligase. Plasmid pPM907 was generated in two steps, by inverse PCR with primers oligoPM434 and oligoPM439 using pStA1CZ as template, DpnI treatment and PCR product circularisation with T4 Ligase, followed by inverse PCR with primers oligoPM519 and oligoPM522, DpnI treatment and PCR product circularisation with T4 Ligase. Plasmid pPM908 was generated in two steps, by inverse PCR with primers oligoPM434 and oligoPM438 using pStA1DZ as template, DpnI treatment and PCR product circularisation with T4 Ligase, followed by inverse PCR with primers oligoPM519 and oligoPM523, DpnI treatment and PCR product circularisation with T4 Ligase. Plasmid pPM909 was generated in two steps, by inverse PCR with primers oligoPM434 and oligoPM439 using pStA1DZ as template, DpnI treatment and PCR product circularisation with T4 Ligase, followed by inverse PCR with primers oligoPM519 and oligoPM523, DpnI treatment and PCR product circularisation with T4 Ligase. Plasmid pPM910 was generated in two steps, by inverse PCR with primers oligoPM434 and oligoPM439 using pStA1EZ as template, DpnI treatment and PCR product circularisation with T4 Ligase, followed by inverse PCR with primers oligoPM519 and oligoPM524, DpnI treatment and PCR product circularisation with T4 Ligase.

**pPM64-lacZ and pPM65-lacZ**
Plasmid pPM64-lacZ was constructed by amplification of the *lacZ* cassette of plasmid pUC19 with primers oligoPM383 and oligoPM384, digestion with NotI and NheI and ligation to vector pMTL-JH16 cut with the same enzymes. Plasmid pPM65-lacZ was generated by insertion of the TT2 terminator from the *C. pasteurianum fdx* gene between the right homology arm and the Gram+ replication origin in pPM64-lacZ by inverse PCR with primers oligoPM563 and oligoPM564, followed by circularisation with KLD Enzyme Mix.

### 1.2. Generation of level 0 plasmids

**pPM911 (FLAG-*gusA*)**
The *gusA* coding sequence from plasmid pPM12 [4] with an additional 30 bp sequence encoding an N-terminal FLAG-tag (MDYKDDDKL) was custom synthesised as gBlock Gene Fragments (IDT) and cloned to pPM900-ccdB StA level 0 plasmid to generate pPM911.

**pPM921 (*adhE2*), pPM924 (*bktb*) and pPM934 (*ter*)**
Coding sequences of *adhE2* (AAK09379), *bktb* (Q0KBP1) and *ter* (WP_002681770) were codon optimised for expression in *C. acetobutylicum* using the Codon Optimization Tool (IDT). Optimised sequences were custom synthesised as gBlock Gene Fragments (IDT) and cloned to pPM900-ccdB StA level 0 plasmid to generate pPM921 (*adhE2*), pPM924 (*bktb*) and pPM934 (*ter*) plasmids.

**pPM926-pPM932 (RBS library)**
Plasmid pStA0-RBS encoding the $RBS_{thl}$ was generated by inverse PCR with primers oligoPM479 and oligoPM480 using plasmid pStA0 as template, followed by circularisation with KLD Enzyme Mix (NEB). The RBS library in level 0 plasmid was generated by inverse PCR with primers oligoPM518 and oligoPM491 (RBS O1, plasmid pPM926), oligoPM518 and oligoPM487 (RBS O2, plasmid pPM927), oligoPM518 and oligoPM492 (RBS O4, plasmid pPM928), oligoPM518 and oligoPM490 (RBS O11, plasmid pPM929), oligoPM518 and oligoPM517 (RBS O20, plasmid pPM930), oligoPM518 and oligoPM489 (RBS O24, plasmid pPM931) and oligoPM518 and oligoPM488 (RBS O28, plasmid pPM932) using pStA0-RBS plasmid as template. PCR products were treated with DpnI and circularised with T4 Ligase.

**pPM957 ($P_{rhaBAD}$)**
$P_{rhaBAD}$ fragment was PCR amplified with primers oligoPM617 and oligoPM618 using plasmid pCK302[3] as template. The PCR product was DpnI treated, the reaction mixture was heat inactivated and used in level 0 reaction with the recipient plasmid pPM900-ccdB to generate pPM957 ($P_{rhaBAD}$).

### 1.3. Generation of level 1 plasmids

**pPM87 plasmid library**
Plasmid library pPM87 (pPM87-O1, O2, -O4, -O11, -O20, -O24, -O28, -R1, -R2, -R3, -R4, -R5 and -R6) was generated by performing individual StA reactions for each plasmid. Each level 1 reaction contained the promoter part ($P_{rhaBAD}$, pPM957), an appropriate RBS part (pPM926-pPM932, pGT330-335), *eyfp* donor plasmid (pGT431) and the recipient plasmid pPM901 (level 1 AB α-δ). Reaction mixtures were transformed to *E. coli*. Cultures originating from single colonies were grown to early exponential phase of growth and analysed with flow cytometry as described before [1].

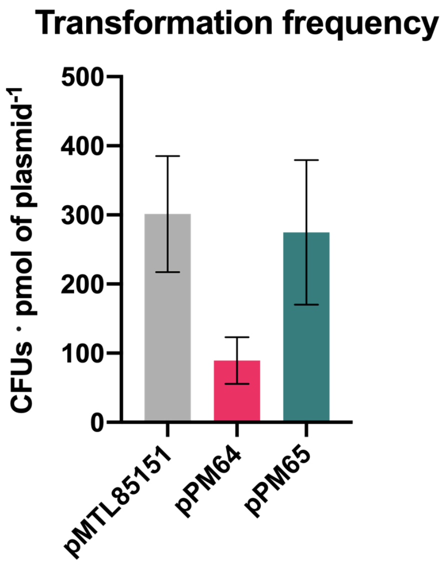


**Figure S1.** Transformation frequencies of plasmids in *C. acetobutylicum* ATCC 824. Error bars represent standard errors of means of three independent experiments.

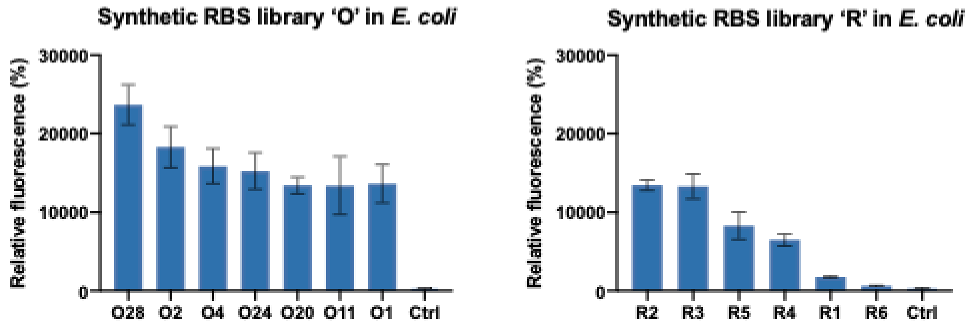


**Figure S2.** Synthetic RBSs tested in *E. coli* DH5α. Level 1 assembly was used to generate plasmid library pPM87, including promoter $P_{rhaBAD}$ (from pPM957), each context-resistant synthetic RBS newly synthesized in the present study O1, O2, O4, O11, O20, O24 and O28 (from pPM926-932) or previously-reported synthetic RBS R1-6 (from pGT330-335), *eyfp* CDS (from pGT431) and vector pPM901 (AB α-δ). Fluorescence of mid-exponential cells growing in LB medium supplemented with 0.33 mg/mL L-rhamnose was measured as described previously [12]. A strain transformed with empty pPM901 was used as a negative control (Ctrl).

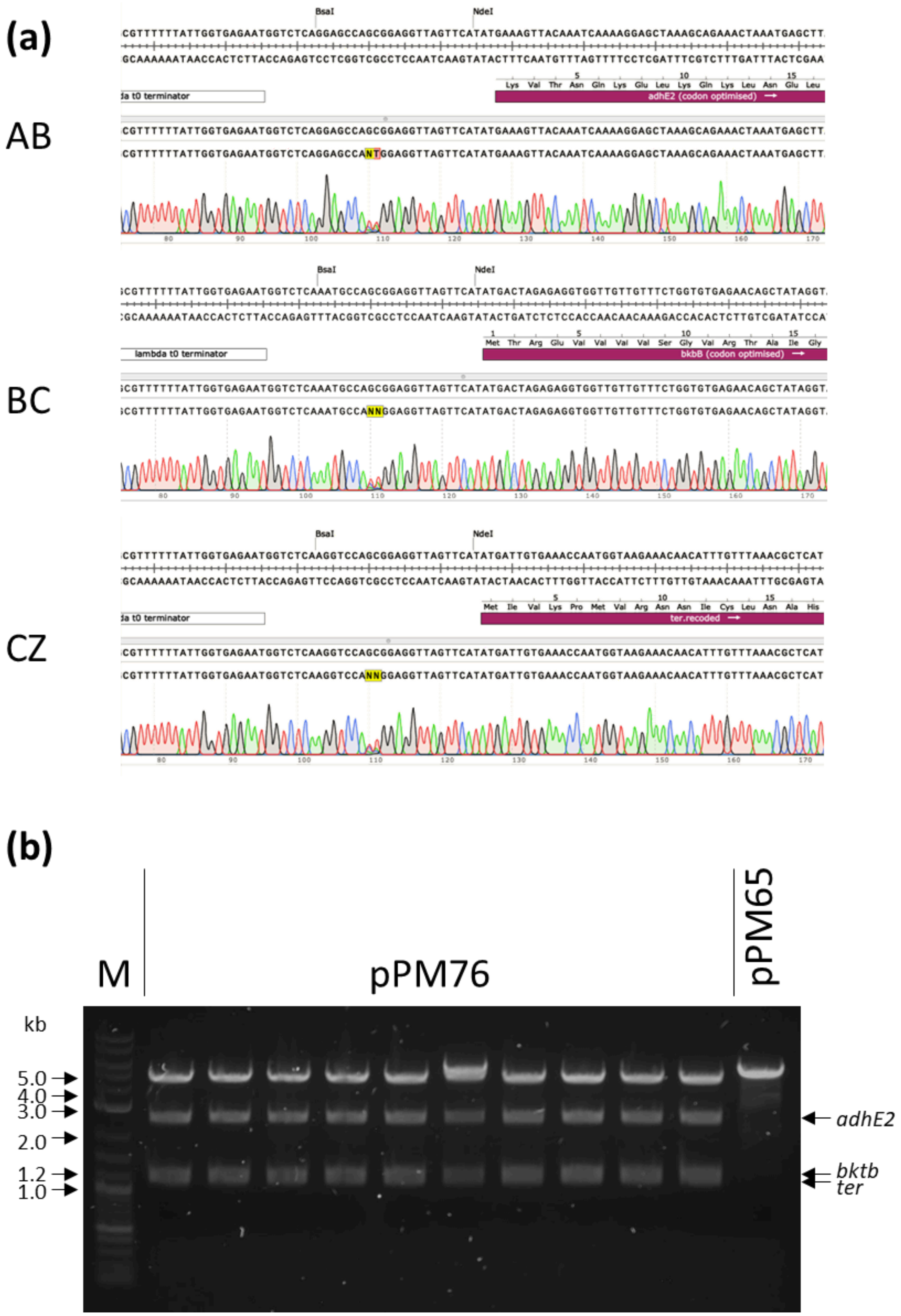


**Figure S3.** Generation of the pPM76 plasmid library encoding the combinatorial extended carbon chain alcohol pathway **(a)** Randomisation of RBSs at the Level 1 plasmids. Plasmid DNA extracted from Level 1 libraries was sequenced using primer oligoPM547. Randomised bases are highlighted in yellow. **(b)** Restriction fragment analysis of 10 random clones from pPM76 plasmid library. Plasmid DNA was purified and digested with enzymes NdeI and NheI (expected fragments 4987, 2600, 1208 and 1200 bp). pPM65 was used as a control (expected fragments 5390 and 53 bp). M - 1 kb Plus DNA Ladder (NEB).

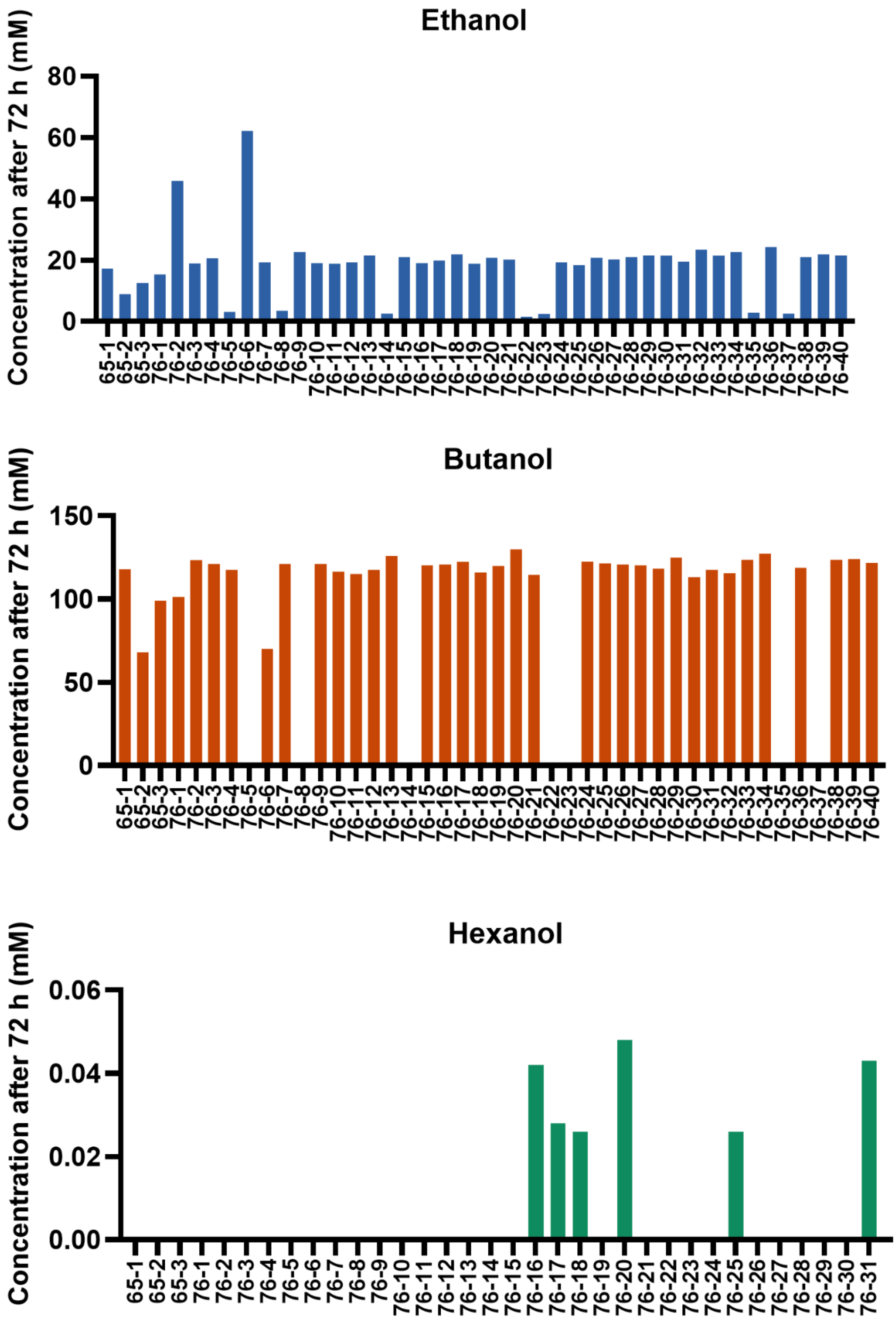


**Figure S4.** Recombinant expression of the combinatorial 1-hexanol pathway. *C. acetobutylicum* was transformed with plasmid library pPM76 encoding *adhE2*, *bktb* and *ter* under synthetic RBSs. A strain harbouring an empty pPM65 plasmid was used as a negative control. Pathways were integrated into the *thl* gene locus. Integrants were selected with erythromycin. Strains were grown in CBMS medium for 72 h. Concentration of ethanol, butanol and 1-hexanol in the culture supernatant was quantified using GC−MS.
